# Loss-of-function *ROX1* mutations suppress the fluconazole susceptibility of *upc2A*Δ mutation in *Candida glabrata*, implicating additional positive regulators of ergosterol biosynthesis

**DOI:** 10.1101/2021.10.07.463606

**Authors:** Tomye L. Ollinger, Bao Vu, Daniel Murante, Josie E. Parker, Lucia Simonicova, Laura Doorley, Mark A. Stamnes, Steven L. Kelly, P. David Rogers, W. Scott Moye-Rowley, Damian J. Krysan

## Abstract

Two of the major classes of antifungal drugs in clinical use target ergosterol biosynthesis. Despite its importance, our understanding of the transcriptional regulation of ergosterol biosynthesis genes in pathogenic fungi is essentially limited to the role of hypoxia and sterol-stress induced transcription factors such as Upc2 and Upc2A as well as homologs of Sterol Response Element Binding (SREB) factors. To identify additional regulators of ergosterol biosynthesis in *Candida glabrata*, an important human fungal pathogen with reduced susceptibility to ergosterol biosynthesis inhibitors relative to other *Candida* spp., we used a serial passaging strategy to isolate suppressors of the fluconazole hypersusceptibility of a *upc2A*Δ deletion mutant. This led to the identification of loss of function mutants in two genes: *ROX1*, the homolog of a hypoxia gene transcriptional suppressor in *Saccharomyces cerevisiae*, and *CST6*, a transcription factor that is involved in the regulation of carbon dioxide response in *C. glabrata*. Here, we describe a detailed analysis of the genetic interaction of *ROX1* and *UPC2A*. In the presence of fluconazole, loss of Rox1 function restores *ERG11* expression to the *upc2A*Δ mutant and inhibits the expression of *ERG3* and *ERG6*, leading to increased levels or ergosterol and decreased levels of the toxic sterol, 14α methyl-ergosta-8,24(28)-dien-3β, 6α-diol, relative to *upc2A*Δ. Our observations establish that Rox1 is a negative regulator of *ERG* gene biosynthesis and indicate that a least one additional positive transcriptional regulator of *ERG* gene biosynthesis must be present in *C. glabrata*.

**Importance:** *Candida glabrata* is one of the most important human fungal pathogens and has reduced susceptibility to azole class inhibitors of ergosterol biosynthesis. Although ergosterol is the target of two of the three classes of antifungal drugs, relatively little is known about the regulation of this critical cellular pathway. Sterols are both essential components of the eukaryotic plasma membrane and potential toxins; therefore, sterol homeostasis is critical for cell function. Here, we identified two new negative regulators of *C. glabrata* of ergosterol (*ERG*) biosynthesis gene expression. Our results also indicate that in addition to Upc2A, the only known activator of *ERG* genes, additional positive regulators of this pathway must exist.s

## Introduction

*Candida* species are a very common cause of human fungal infections. Candidiasis encompasses infections of mucosal and epidermal tissues such as oropharyngeal candidiasis and vulvovaginal candidiasis as well as invasive infections of deep organs and the bloodstream (1). Mucosal infections affect both immunocompetent and immunocompromised people while invasive infections primarily affect critically ill patients or those with altered immune function. Although *Candida albicans* is the most prevalent infecting species, the so-called non-albicans *Candida* spp. such as *C. glabrata, C. parapsilosis* (2) and, most recently, *C. auris* have emerged as clinically significant causes of candidiasis (3). Of these, *C. glabrata* is the second most common cause candidiasis in most clinical case series of invasive infections (1,2).

One of the most important characteristics that distinguishes *C. glabrata* from *C. albicans* is that *C. glabrata* is less susceptible to azole antifungal drugs and has the highest rate of resistance to echinocandins; thus, *C. glabrata* has reduced susceptibility to two of the three classes of drugs currently used to treat invasive fungal infections (4). The rate of azole and echinocandin resistance varies significantly by institution (5). For example, some institutions report almost no resistance while others have fluconazole resistance rates as high as 18%. In contrast, the rates of *C. albicans* azole resistance remains uniformly quite low. Genomic sequences of azole-resistant *C. albicans* isolates indicate that mutations in multiple genes can lead to resistance including gain-of-function mutations in transcription factors regulating drug efflux pump expression, mutations in the azole target *ERG11*, and gain-of-function mutations in *UPC2*, a transcription factor that regulates the expression of ergosterol biosynthesis genes during periods of sterol depletion (6). Curiously, mutations associated with azole resistance in *C. glabrata* clinical isolates are essentially limited to gain-of-function alleles in *PDR1*, the transcription factor that regulates efflux pump expression, suggesting that these two species adapt to azole-induced sterol depletion in very different ways (7, 8).

Although gain of function mutations in the *C. glabrata* ortholog of *C. albicans* Upc2, *UPC2A* have not been isolated in fluconazole-resistant *C. glabrata* clinical isolates, Whaley et al. showed that *UPC2A* was essential for fluconazole resistance in *PDR1* gain-of-function mutants (9). In the absence of sterol stress such as hypoxia or azole exposure, *UPC2A* has a modest effect on ergosterol gene (*ERG*) expression (9, 10, 11); however, under these inducers of sterol stress, *upc2A*Δ mutants show a dramatic reduction in fitness (9, 10). Previous studies in *C. glabrata* indicate that Upc2A regulates genes involved in the uptake of exogenous sterols during hypoxic growth (11). However, little is known about how *UPC2A* interacts with other genes involved ergosterol biosynthesis or the cellular response to sterol depletion. In *S. cerevisiae*, the Upc2A homolog *Sc*Upc2 functions with *Sc*Ecm22 to regulate sterol biosynthesis (12). The *upc2*Δ *ecm22*Δ double mutant is viable but hypersusceptible to inhibitors of the ergosterol pathway and resistant to amphotericin B (12). In *C. glabrata*, Upc2B is homologous to Upc2A but *upc2B*Δ mutants do not show increased susceptibility to fluconazole or decreased growth under hypoxic conditions (10). These observations suggest that the transcriptional regulation of the ergosterol pathway in *C. glabrata* is distinct from the model yeast *S. cerevisiae*.

More generally, the regulation of *ERG* gene expression in fungal pathogens is remarkably understudied given the importance of the ergosterol biosynthesis pathway to the treatment of fungal infections (13); two of the three main therapies, azoles and polyenes, target ergosterol. To date, the main focus of this area of research has been on factors related to mammalian Sterol Response Element Binding (SREB) proteins or the Zinc(2)-Cys(6) factors such as Upc2A that function similarly to SREB factors but have no sequence homology to those proteins (14). Furthermore, if we consider the fact that these factors are not essential and have little effect on ergosterol levels or growth in the absence of sterol stress, then additional pathways and factors must regulate the expression of *ERG* genes.

To identify additional regulators of the *ERG* pathway under sterol stress, we took advantage of the fact that deletion of *UPC2A* prevents increased efflux pump expression from causing azole resistance and designed a laboratory evolution experiment for the isolation of suppressors of *upc2A*Δ fluconazole hyper-susceptibility (9). In wild type cells, an experimental laboratory evolution approach in the presence of fluconazole would be expected to lead to gain-of-function mutants in Pdr1, the TF that regulates the expression of efflux pumps *CDR1/2*. We hypothesized that the presence of a *upc2A*Δ mutation would prevent the emergence of *PDR1* gain-of-function mutations and allow us to identify either gain-of-function mutants in other positive regulators of *ERG* gene expression or, alternatively, loss of function mutants in negative regulators.

As we described below, this laboratory evolution strategy led to the isolation loss-of-function mutations in two transcription regulators: *ROX1* and *CST6. ROX1* is homologous to a repressor of hypoxia genes in *S. cerevisiae* (15) while *CST6* is a homolog of a *S. cerevisiae* and a *C. albicans* (Rca1) transcriptional regulator of carbonic anhydrase expression (16). In this report, we focused our characterization on the transcriptional and biochemical changes that mediate the suppression of *upc2A*Δ fluconazole hyper-susceptibility by *ROX1* loss of function mutations; a detailed analysis of the interaction of *CST6* with *UPC2A* awaits additional work.

## Results

### Experimental laboratory evolution generates suppressors of *upc2A*Δ fluconazole hypersusceptibility

The general strategy for our experimental laboratory evolution experiment is outlined in Fig. 1A. First, we constructed a *upc2A*Δ mutant in the His^-^, BG2 genetic background (17) using a recyclable dominant selectable marker system recently reported by members of our team (18). Three founder colonies were selected and 88 lineages from each were inoculated into a microtiter plate containing YPD medium supplemented with fluconazole at 1/8 MIC (0.25 µg/mL) of the parental strains (Fig. 1). The cultures were grown to stationary phase and then passaged to a 2-fold higher concentration of fluconazole until a final concentration of 64 µg/mL (32X MIC) was reached. All but 20 lineages went extinct before reaching a fluconazole concentration of 64 µg/mL. The contents of wells that grew at 64 µg/mL were plated on YPD as well as on synthetic medium lacking histidine to eliminate possible environmental contaminants. Inspection of the YPD plates indicated that the majority of wells contained heterogenous mixtures of small colonies and rarer large colonies, although some wells had large colonies only (Fig. 1B).

**Figure 1.**
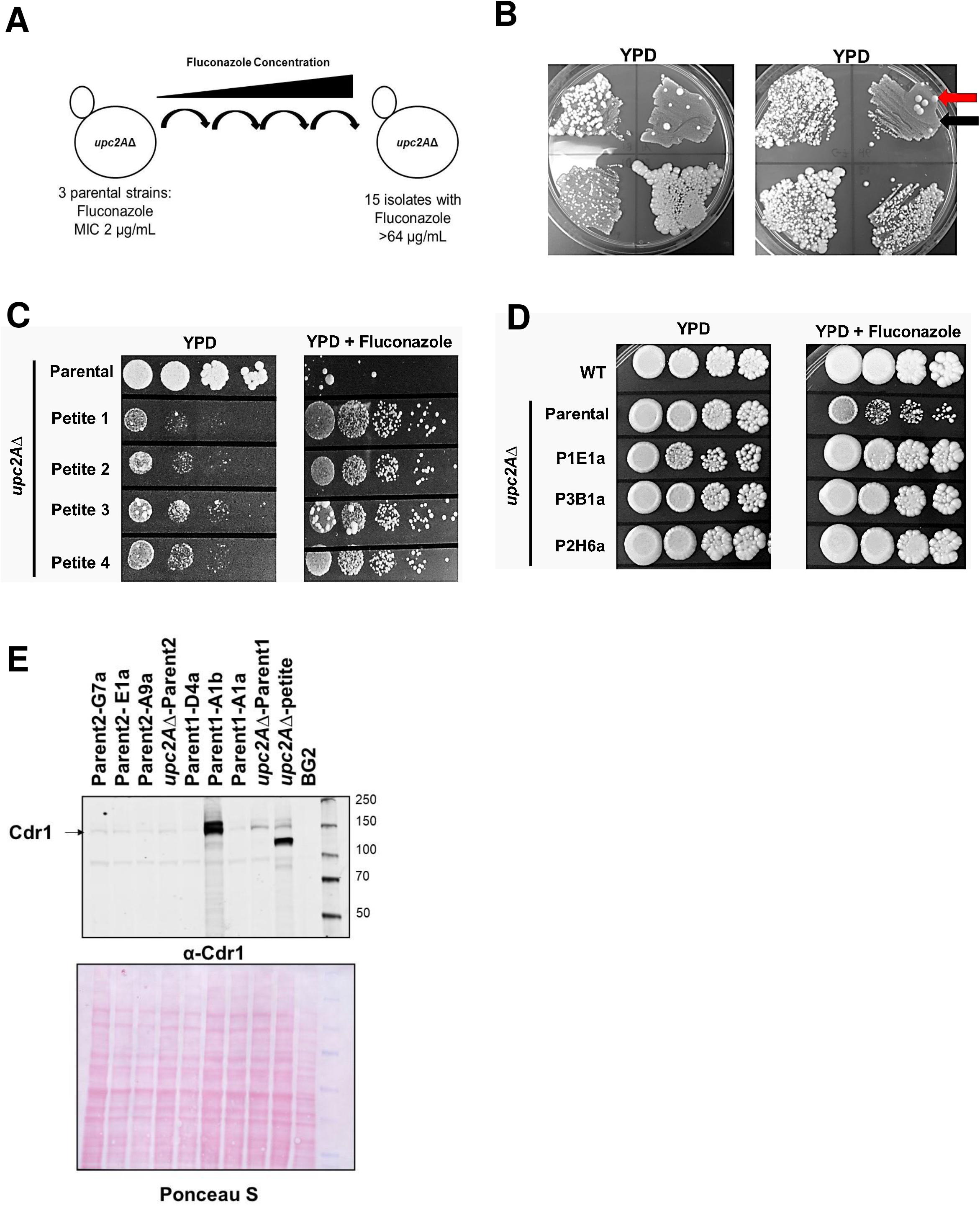
Experimental laboratory evolution identifies suppressors of *upc2A*Δ fluconazole hyper-susceptibility. A. Schematic of selection strategy. B. Representative plates inoculated with the contents of wells with growth at 64 µg/mL of fluconazole. The red arrow indicates large colony isolates and the red arrow indicates petite colony isolates. C. Spot dilution assays of representative petite colony isolates on YPD and YPD + 40 µg/mL fluconazole. D. Spot dilution assays of representative large colony isolates on YPD and YPD + 40 µg/mL fluconazole. E. Western blot analysis of representative large colony and petite isolates for Cdr1 after growth in YPD to log phase. The blot is representative of two biological replicates which showed the same pattern.

We purified both types of colonies by serially passaging on YPD plates without fluconazole three times. The small colony isolates retained this colony morphology in the absence of selective pressure. These small colonies also grew poorly on glucose relative to the parental *upc2A*Δ strains but were much more fit than the parental *upc2A*Δ in the presence of fluconazole (Fig. 1C). The small colony size and poor growth on glucose strongly suggested that these strains might have mitochondrial defects (19). Consistent with that hypothesis, the small colony isolates were unable to grow on glycerol plates and showed decreased mitochondrial DNA staining (data not shown), consistent with petite isolates. Previous work has shown that petite *C. glabrata* strains as well as those with other mitochondrial defects have increased Pdr1 activity, elevated expression of the Cdr1 efflux pump and reduced fluconazole susceptibility (19).

From the large colony isolates, 11 retained decreased fluconazole susceptibility relative to the parental *upc2A*Δ strains after serial passaging on YPD plates (Fig. 1D). The growth curves for two representative resistant isolates are provided in Fig. S1, further confirming the resistance of the strains to fluconazole relative to the *upc2A*Δ. As discussed above, the most common mechanism of fluconazole resistance is Pdr1 gain-of-function mutations that lead to increased expression of Cdr1, a putative fluconazole efflux pump (7, 8). To determine if any of the 11 large colony isolates had increased Cdr1 expression, we analyzed the protein levels of Cdr1 protein in the suppressor strains by western blot (Fig. 1E). Of the 11 suppressors, only two strains showed evidence of increased Cdr1 expression relative to the parental; the petite strain and P1A1b, a non-petite mutant. Thus, the decreased fluconazole susceptibility for the majority (10/11) of the suppressor mutants cannot be explained by Pdr1-mediated efflux pump activation. As noted above, petite strains of *C. glabrata* activate Pdr1 and these data further support the assignment of these strains as petites (19). Previous experiments in a clinical *C. glabrata* isolate indicated that *upc2A*Δ strongly reduced the effect of Pdr1 gain-of-function on fluconazole susceptibility (9). Our observations, however, indicate that under some circumstances Pdr1 activation can lead to Upc2A independent reduction in fluconazole susceptibility but *upc2A*Δ mutation largely prevents the emergence of *PDR1* gain-of-function mutations as a mechanism of fluconazole resistance.

### Genomic sequencing identifies putative loss of function in mutations in *ROX1* and *CST6* in the majority of *upc2A*Δ fluconazole suppressor strains

The parental *upc2A*Δ founder strains as well as 12 suppressor strains were resequenced. Following analysis using the Lasergene suite of software, 11 strains were found to contain mutations within coding regions (Fig. 2A). Of these 11, 6 contained either frameshift or nonsense mutations that that would lead to loss of function in CAGL0I05170g (17, 20), a homolog of *S. cerevisiae* transcription factor Sc*CST6* (*C. glabrata* gene and protein names will not have prefix modifiers while *C. albicans* and *S. cerevisiae* genes and protein names will have species prefixes). Four strains contained similar types of loss-of-function mutations in CAGL0D05434g, homologous to the *S. cerevisiae* High Mobility Group (HMG) transcriptional repressor *ROX1* (15, 17). One strain contained a nonsense codon in CAGL0D03828g, an uncharacterized homolog of the *S. cerevisiae* mediator component, *MED6*; this is predicted to be an essential protein and was not pursued further. One *rox1* and one *cst6* mutant strain also had additional non-synonymous polymorphism in the cell surface adhesion *PWP3*. Finally, a *cst6* mutant strain also had a non-synonymous polymorphism in the putative glucose transporter *HXT4*.

**Figure 2.**
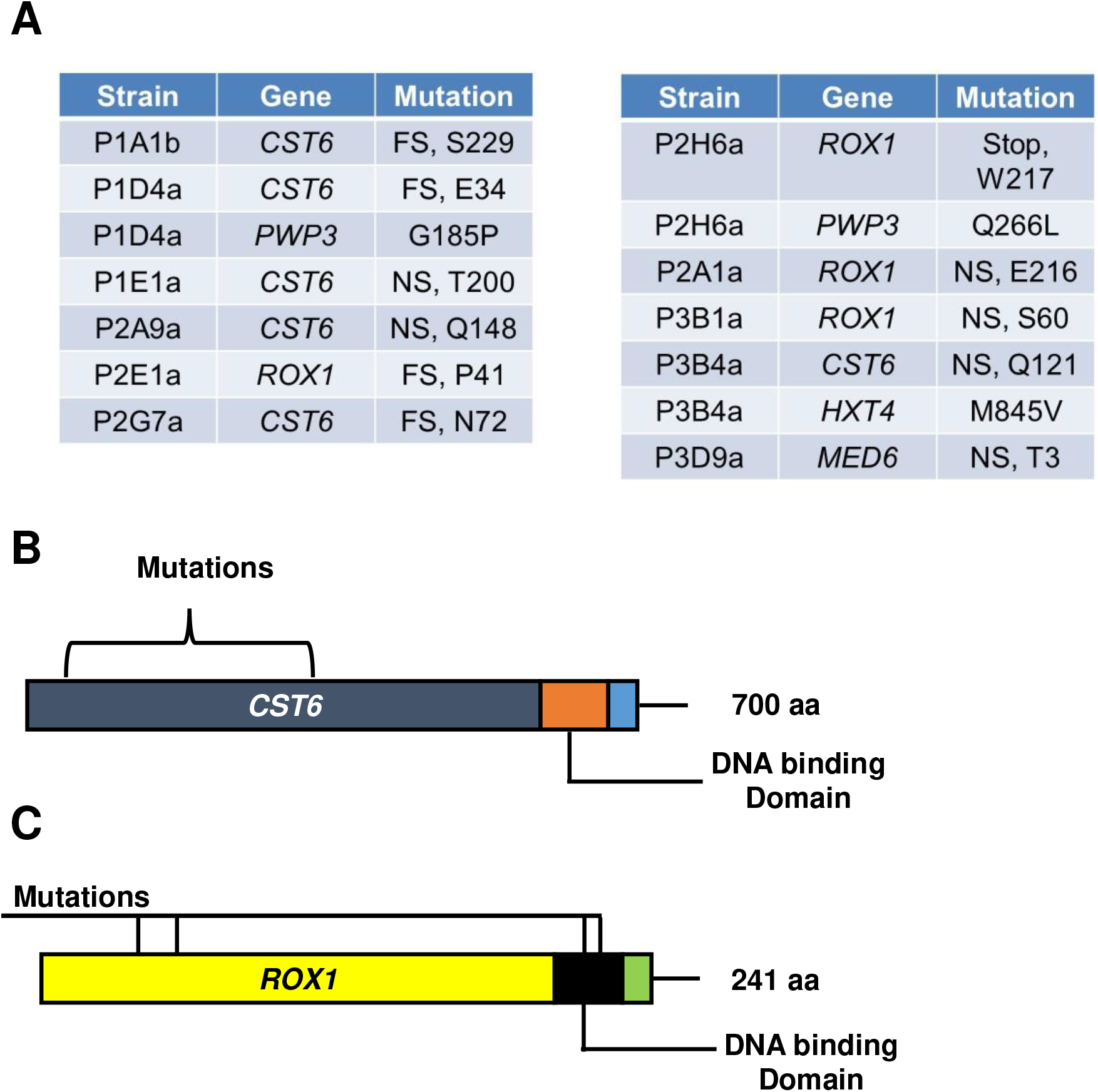
Loss of function mutations in *ROX1* and *CST6* are present in the *upc2A*Δ fluconazole-suppressor strains. A. The table indicates the code for the individual isolate and the coding region mutation identified in that isolate. B. Location of mutations in *CST6*. Location of mutations in *ROX1*.

In *S. cerevisiae, Sc*Cst6 has been shown to regulate the expression of carbonic anhydrase (*NCE103*) and deletion mutants of *CST6* have similarly been shown to play a modest role in the expression of *NCE103* and CO_2_ sensing in *C. glabrata* (21). In *C. albicans, CaRCA1* is the homolog of Sc*CST6* and *CST6*. Ca*RCA1* plays a role in *NCE102* expression (21) but also appears to affect ergosterol biosynthesis as evidenced by the observations that Ca*rca1*ΔΔ mutants have decreased susceptibility to fluconazole and reduced levels of ergosterol relative to WT (22). A library of *C. glabrata* deletion mutants contains a *cst6*Δ strain but it was not reported to have altered susceptibility to fluconazole. The putative *CST6* loss of function mutations identified in these strains are frameshift or nonsense mutations within the N-terminal third of the protein, well short of the basic leucine zipper (bZip) DNA binding domain (Fig. 2B).

In *S. cerevisiae*, Rox1 is a well-studied HMG repressor of hypoxia-related genes including ergosterol biosynthesis genes. Indeed, *Sc*Rox1 functions to repress some genes that are activated by *Sc*Upc2 and its ortholog *Sc*Ecm33 in response to either ergosterol depletion or reduced oxygen availability (23). For example, previous work from the Edlind lab has shown that deletion of *ScROX1* decreases the susceptibility of *S. cerevisiae* to azole antifungals 4-to 10-fold and increases the expression of *ERG* genes (24). However, the role of *ROX1* in *C. glabrata* has not been investigated previously besides as part of the large-scale library screening (17). This screen found that *rox1*Δ was more susceptible to amphotericin but similar to wild type in the presence of fluconazole. Interestingly, although both *Sc*Rox1 and Rox1 contain homologous DNA binding domains, the HMG DNA binding domain for ScRox1 is positioned at the N-terminus, while the DNA binding domain of Rox1 is at the C-terminus (Fig. 2C). The *ROX1* mutations are either nonsense mutations early in the N-terminal portion or within the DNA binding domain (Fig. 2C).

### Loss of function mutations of *ROX1* suppress fluconazole hyper-susceptibility of *upc2A*Δ strains

To confirm that the loss of function mutations identified in the *upc2A*Δ fluconazole suppressor strains were responsible for this phenotype, we attempted to make the corresponding double mutant deletion strains. In the case of *ROX1*, we were able to generate the *rox1*Δ and *rox1*Δ *upc2A*Δ strains in the KKY2001 background; in addition, we generated a *rox1*Δ mutation in a strain lacking *PDR1*, a transcription factor that has a profound effect on fluconazole susceptibility. In the presence of fluconazole, *upc2A*Δ and *pdr1*Δ cells have a significant growth defects relative to WT while *rox1*Δ is slightly less susceptible at high concentrations of fluconazole after 24 hr but not at 48 hr (Fig. 3A). Deletion of *ROX1* in the *pdr1*Δ background nearly restores WT growth at the lowest concentrations of fluconazole after 48 hrs but has minimal effect on the increased fluconazole susceptibility of the *pdr1*Δ mutant at higher concentrations. In contrast, deletion of *ROX1* in the *upc2A*Δ mutant restores growth to near wild type rate at the lower concentrations of fluconazole and at longer incubation times. This strongly supports the conclusion that the loss of function mutations in *ROX1* identified in the evolved strains are responsible for the reduced fluconazole susceptibility relative to the *upc2A*Δ parental strains. In addition, these data indicate that loss of Rox1 function is not a global suppressor of fluconazole susceptibility since the mutation has minimal effects on WT or *pdr1*Δ cells.

**Figure 3.**
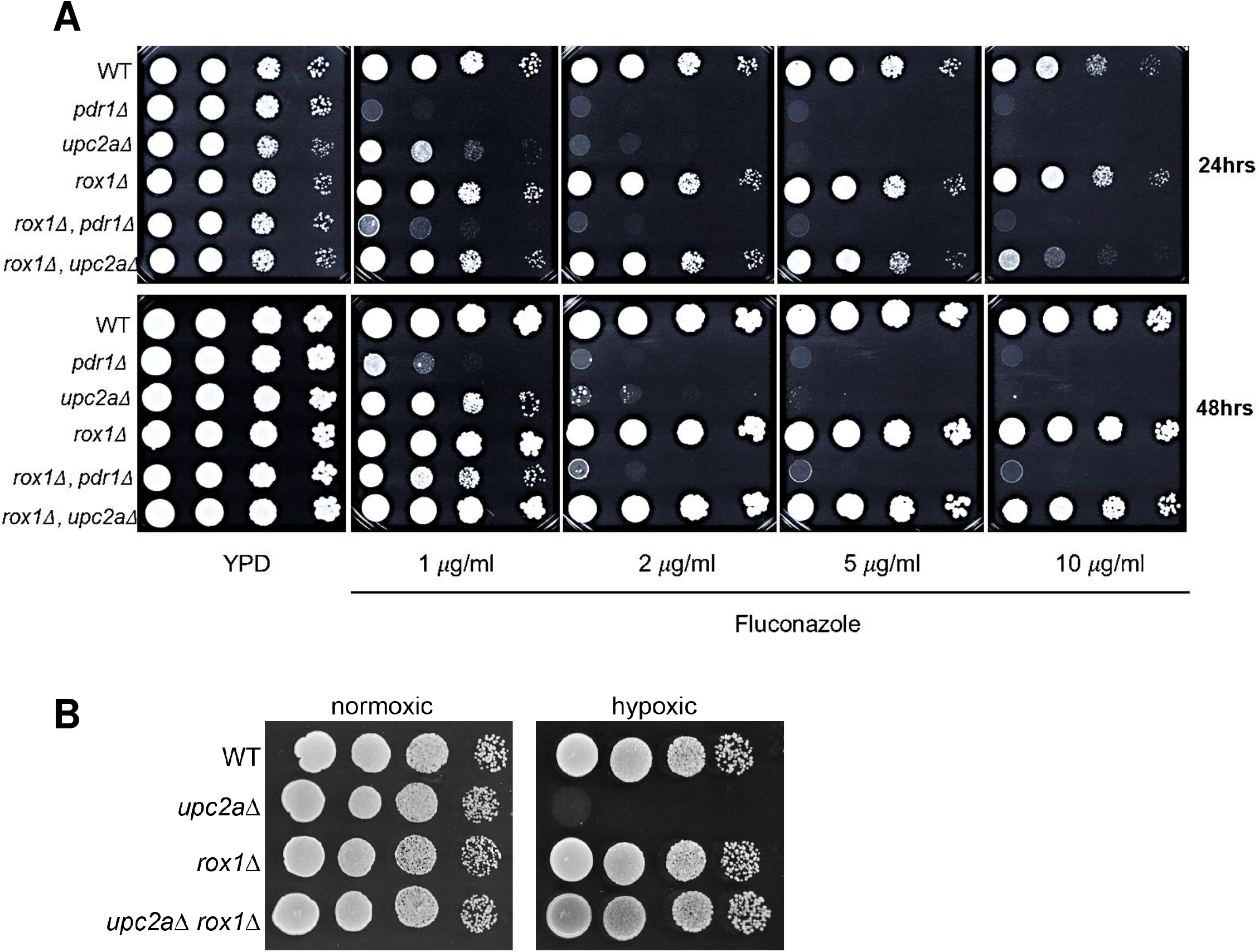
Deletion of *ROX1* suppresses fluconazole and hypoxia hyper-susceptibility of *upc2A*Δ mutants. A. Ten-fold dilution series of the indicated strains were spotted on YPD or YPD plates with the indicated concentrations of fluconazole and incubated for 24 or 48 hr at 30°C. B. The indicated strains were plated on YPD and incubated at 30°C in ambient air or in a GasPak to establish a low oxygen environment as described in Materials and Methods.

Despite multiple attempts to generate a *cst6*Δ *upc2A*Δ double mutant using both standard recombination and CRISPR/Cas9 approaches, we were unable to do so. The *cst6*Δ deletion mutant was generated as part of the large library of *C. glabrata* deletion mutants. In addition, others have reported the construction of *cst6*Δ mutants in other contexts (20). We obtained the *cst6*Δ strain from the library collection and determined its susceptibility to fluconazole; in CLSI microdilution assays, it is less susceptible to fluconazole than the parental strain (MIC *cst6*Δ: 16 µg/mL vs. WT: 4 µg/mL). We elected to focus the remained of our characterization of the interaction between Rox1 and Upc2A since we were able to confirm that loss of function of *ROX1* suppresses the *upc2A*Δ fluconazole hypersusceptibility phenotype.

Upc2A is also required for *C. glabrata* to grow under hypoxic conditions (10, 11) and *Sc*Rox1 was originally identified as a repressor of hypoxia-related genes during growth under aerobic conditions (23). Accordingly, many genes repressed by *Sc*Rox1 during normoxia are also activated by *Sc*Upc2. Therefore, we compared the growth of the *upc2A*Δ *rox1*Δ strain to *upc2A*Δ and *rox1*Δ under low oxygen conditions (Fig. 3B). As expected, the *upc2A*Δ mutant grew poorly while the *rox1*Δ mutant grew similar to wild type. Consistent with its fluconazole phenotype, the *rox1*Δ *upc2A*Δ mutant grew similarly to wild type indicating that *ROX1* loss of function suppressed both the fluconazole and hypoxia susceptibility of *upc2A*Δ.

### Deletion of *ROX1* increases ergosterol content and alters the sterol profile of *upc2A*Δ mutants

To gain insights in to how the *rox1*Δ affects the sterol content and distribution (Fig. 4) in the presence and absence of fluconazole, we used mass spectrometry to characterize the sterol profile of WT, *upc2A*Δ, *rox1*Δ, and *rox1*Δ *upc2A*Δ strains in YPD medium with and without 8 µg/mL fluconazole. In the absence of fluconazole, WT, *rox1*Δ and *upc2A*Δ strains had statistically indistinguishable amounts of total cell ergosterol (Fig. 5A) while *rox1*Δ *upc2A*Δ had ∼1.5-fold more ergosterol than the other strains. In the presence of fluconazole concentrations that allowed *upc2A*Δ to grow, albeit at a reduced rate (Supplementary Figure 1), the *rox1*Δ strain had modestly increased ergosterol while the ergosterol content of the *upc2A*Δ mutant was reduced slightly (Fig. 5B). The double mutant *rox1*Δ *upc2A*Δ, in contrast, has nearly 50% more ergosterol than the *upc2A*Δ deletion mutant and had statistically significantly increased ergosterol content than WT (p < 0.05). Interestingly, the proportion of ergosterol within the total cell sterols is reduced in the *rox1*Δ *upc2A*Δ mutant relative to WT while the others are not affected, indicating that the relative distribution of sterols is also altered in this mutant (Fig. 5C).

**Figure 4.**
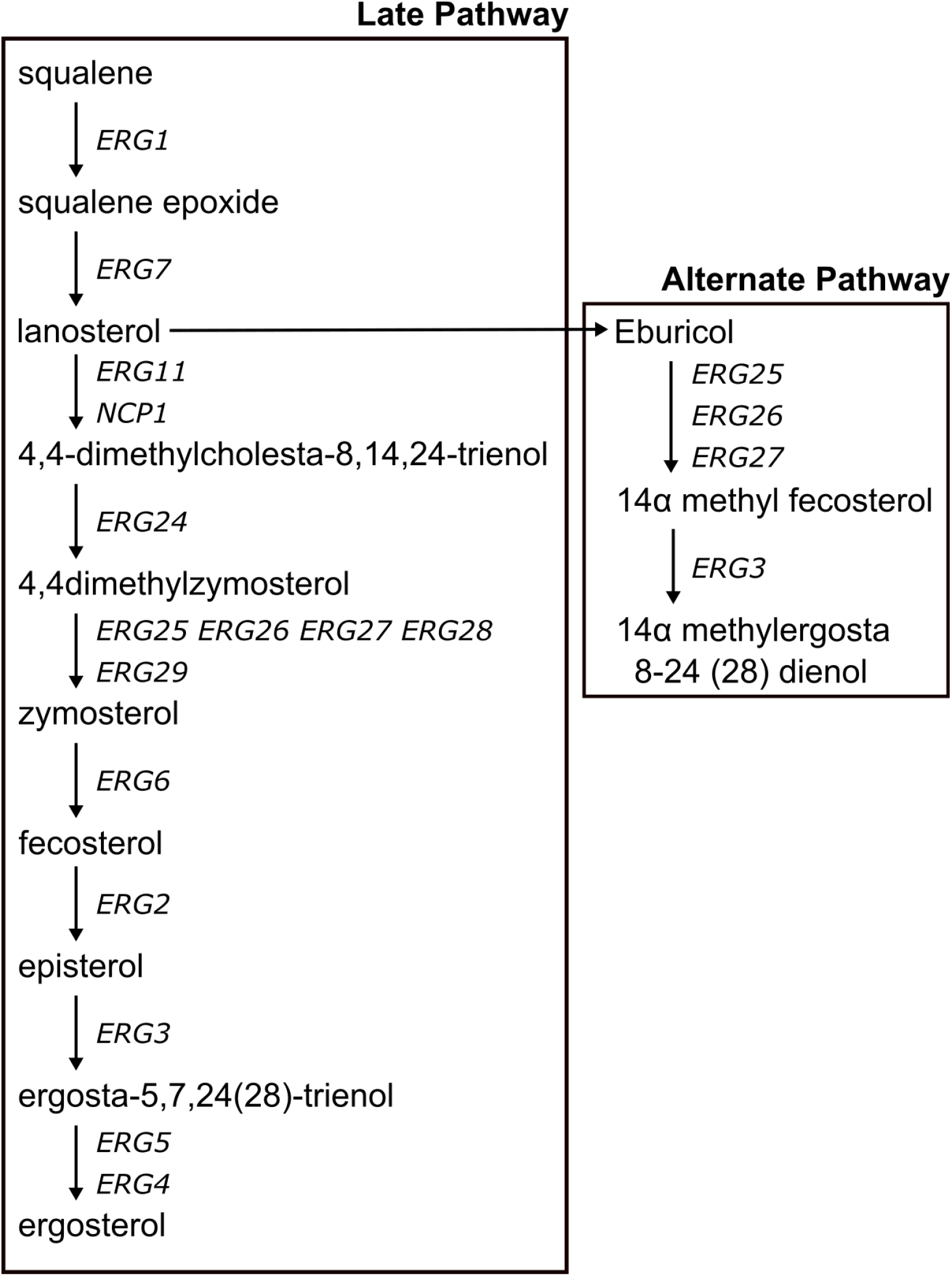
Schematic of the late portion of the ergosterol biosynthesis pathway beginning with squalene.

**Figure 5.**
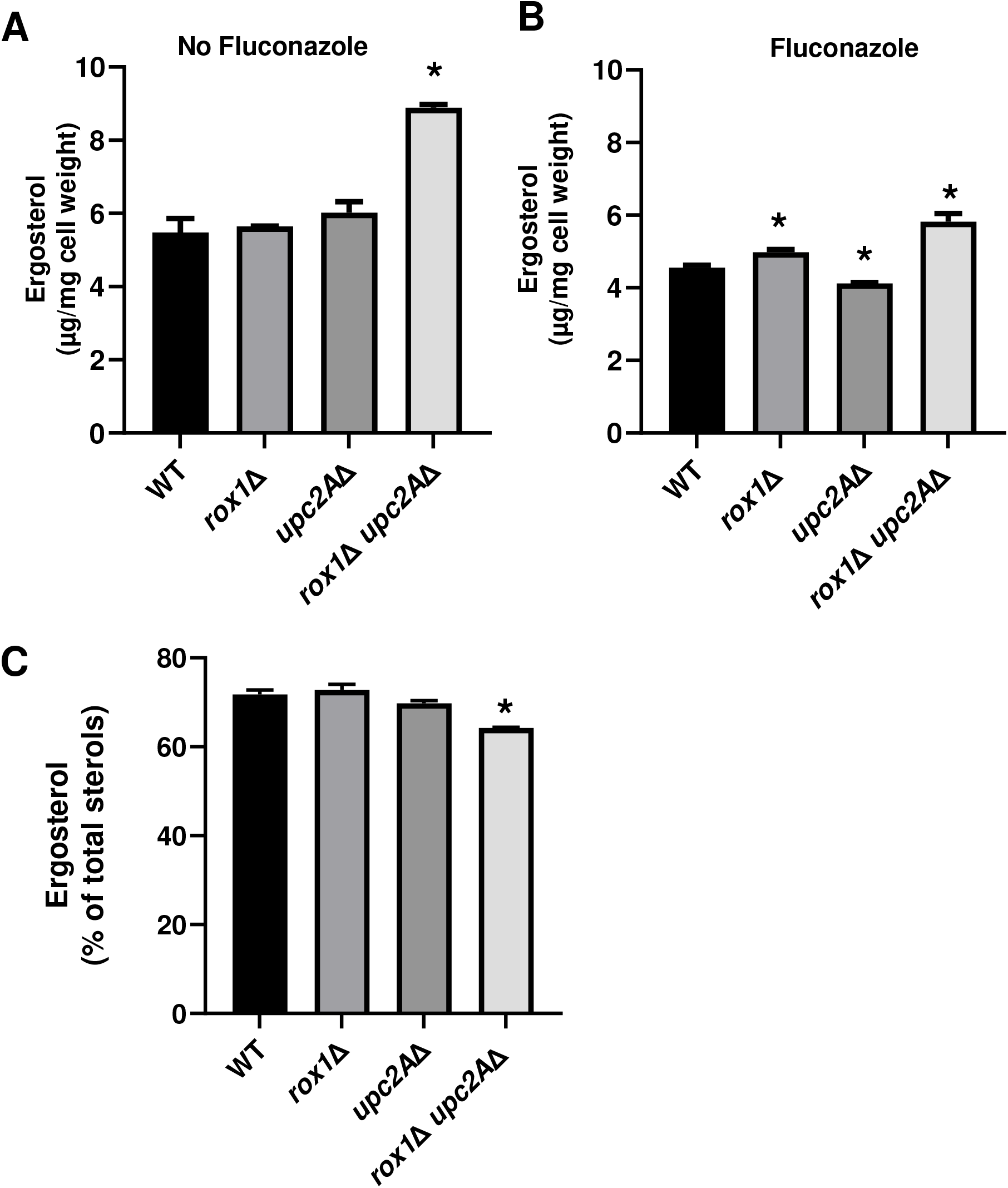
The *rox1*Δ *upc2A*Δ mutant has elevated ergosterol relative to *upc2A*Δ and wild type in the presence and absence of fluconazole. The content of ergosterol was determined as described in Materials and methods from log phase cultures of the indicated strains in YPD without (A) and with fluconazole (8 µg/mL). C. The proportion ergosterol compared to total sterols for the indicated strains was determined under the same conditions as for A and B. The bars indicate the means with standard error of the means. Statistical significance relative wild type is indicated by an *. Data were analyzed by ANOVA followed by Student t tests for individual comparisons with p < 0.05 indicating statistical significance. The complete sterol data are provided in Supplementary Table 1.

The ergosterol biosynthesis intermediate (Fig. 4) that is most altered in the *upc2A*Δ *rox1*Δ mutant in both the presence and absence of fluconazole is zymosterol, a substrate for the C-24 methyltransferase enzyme Erg6. Zymosterol accumulates in both *rox1*Δ and *upc2A*Δ *rox1*Δ strains in the absence of fluconazole but does so only in the double mutant in the presence of fluconazole (Fig. 6A&B). These data suggest that Erg6 activity is reduced in strains lacking lacking *ROX1* and particularly in fluconazole-treated *rox1*Δ *upc2A*Δ mutants. Erg6 also catalyzes the conversion of lanosterol, the substrate of Erg11, into eburicol, the first sterol in the alternate pathway (Fig. 4). Lanosterol is a minor component of total sterols in the absence of fluconazole (WT/YPD: 1.28%, Table S1) but increases dramatically in fluconazole-treated cells (WT/FLC: 20%, Table S1) because the activity of Erg11 is inhibited. As shown in Fig. 6C, lanosterol is further increased in fluconazole-treated *rox1*Δ *upc2A*Δ mutants relative to fluconazole-treated WT. Since ergosterol is not depleted in the *rox1*Δ *upc2A*Δ mutant, the increase in lanosterol seems unlikely to be due to differences in Erg11 activity but rather due to reduced Erg6-mediated conversion of lanosterol to eburicol. Further supporting this notion, eburicol is significantly reduced in fluconazole-treated *rox1*Δ *upc2A*Δ cells relative to WT (Fig. 6D). These data strongly support the idea that the combination of *upc2A*Δ and *rox1*Δ mutations lead to reduced Erg6 activity in the presence of fluconazole.

**Figure 6.**
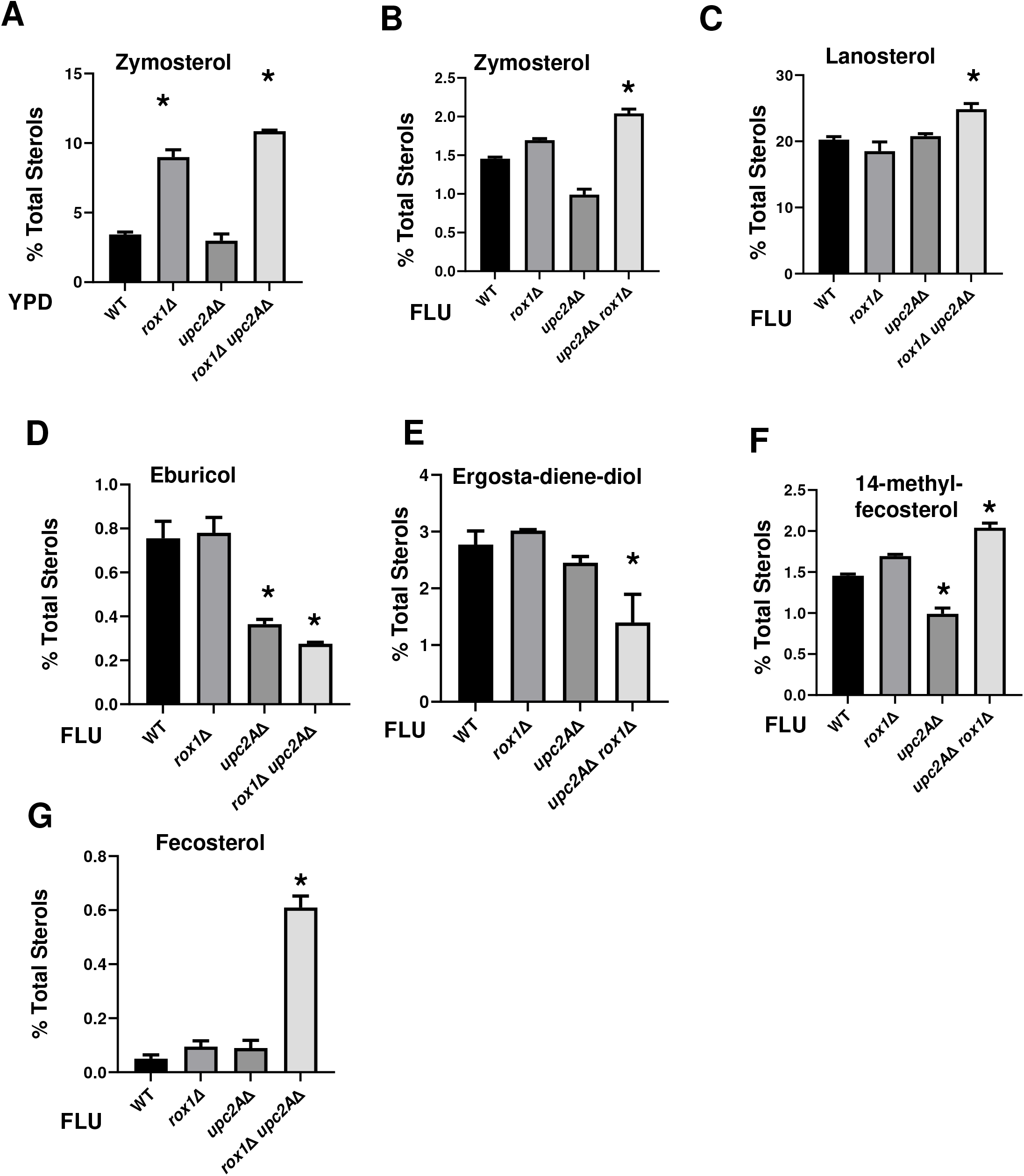
The *rox1*Δ *upc2A*Δ mutant have altered sterol profiles including reduced proportion of the ergosta-diene-diol that mediates fluconazole toxicity. Each of the panels depict the proportion of the indicated sterols determined as described in Materials and methods. A. Zymosterol, YPD. B. Zymosterol, YPD+FLU. C. Lanosterol, YPD+FLU. D. Eburicol, YPD+FLU. E. Ergosta-diene-diol is a shortened name for: 14α methyl-ergosta-8,24(28)-dien-3β, 6α-diol, YPD+FLU. F. 14-Methyl fecosterol, YPD+FLU. G. Fecosterol, YPD+FLU. The bars indicate the means with standard error of the means. Statistical significance relative wild type is indicated by an *. Data were analyzed by ANOVA followed by Student t tests for individual comparisons with p < 0.05 indicating statistical significance. The complete sterol data are provided in Supplementary Table 1.

Ultimately, the alternate pathway converts lanosterol to 14α methyl-ergosta-8,24(28)-dien-3β, 6α-diol (Fig. 4) which is toxic to cells and is thought to be part of the mechanism by which azole drugs inhibit fungal growth (25). In *rox1*Δ *upc2A*Δ mutants treated with fluconazole, the toxic dienol is reduced relative to *upc2*AΔ and WT (Fig. 6E). In addition, the proportion of 14α methyl fecosterol, the immediate precursor to 14α methyl-ergosta-8,24(28)-dien-3β, 6α-diol, is also increased in *rox1*Δ *upc2A*Δ mutants treated with fluconazole (Fig. 6F). Erg3 catalyzes this conversion and the accumulation of 14α methyl fecosterol in the fluconazole-treated *upc2A*Δ *rox1*Δ double suggests that Erg3 activity may also be reduced in that strain.

In the ergosterol pathway, Erg3 also converts episterol to ergosta-5,7,24(28)-trienol. Although episterol is elevated in the fluconazole-treated *rox1*Δ *upc2A*Δ mutant, the variation between samples is too great to confidently conclude that it accumulates. The previous sterol in the pathway is fecosterol and its levels are increased five-fold in the fluconazole-treated *rox1*Δ *upc2A*Δ mutant relative to fluconazole-treated wild type (Fig. 6G). Because Erg3 mediates the conversion of fecosterol to episterol, it is likely that its reduced activity leads to the accumulation of fecosterol as well. Taken together, these data indicate that the *rox1*Δ *upc2A*Δ mutant double mutant appears to have both increased overall ergosterol and reduced accumulation of the toxic diol relative to *upc2*AΔ in the presence of fluconazole. The sterol profiles suggest that the near WT susceptibility of *upc2AΔ rox1Δ* to fluconazole may be due to alterations in the activity of Erg6 and Erg3. Since both Rox1 and Upc2A are transcriptional regulators of *ERG* genes, it seems likely the mechanism of this remodeling of the *rox1*Δ *upc2A*Δ strains sterol composition is due, at least in part, to differential expression of genes such as *ERG3* and *ERG6*.

### *ERG11* gene expression is maintained at WT levels in *rox1*Δ *upc2*AΔ while *ERG3* and *ERG6* expression is reduced in the presence of fluconazole

We hypothesized that altered expression of *ERG* genes such as *ERG3, ERG6* and *ERG11* might be responsible for the changes in sterol content in *rox1*Δ *upc2A*Δ mutants exposed to fluconazole. To test that hypothesis, we determined the expression of *ERG1*-*11*&*24* in WT and *rox1*Δ *upc2A*Δ in the presence and absence of fluconazole using quantitative RT-PCR (Fig. 7A). The first notable feature of the effect of the *rox1*Δ *upc2A*Δ mutant on *ERG* gene expression is that it varies with the specific *ERG* gene. In the absence of fluconazole, for example, *ERG11* expression is elevated 3.5-fold relative in the *rox1*Δ *upc2A*Δ mutant relative to WT (Fig. 7A&B). Since this strain lacks Upc2A, the only known regulator of *ERG* gene expression in *Candida*, the elevated expression of *ERG11* indicates that at least one other positive transcriptional regulator must exist. In the presence of fluconazole, *ERG11* expression in the *rox1*Δ *upc2A*Δ is nearly identical to WT but reduced somewhat relative to the untreated strain, indicating that loss of Upc2A function has an effect on the expression of *ERG11*. Western blot analysis (Fig. 7C) confirmed the RNA levels as shown in Fig. 7B. Thus, the double mutant restores the expression of the fluconazole target Erg11 in the *upc2A*Δ background to that similar to wild type, providing at least a partial explanation for the ability of *rox1*Δ deletion to suppress the *upc2A*Δ fluconazole hypersusceptibility.

**Figure 7.**
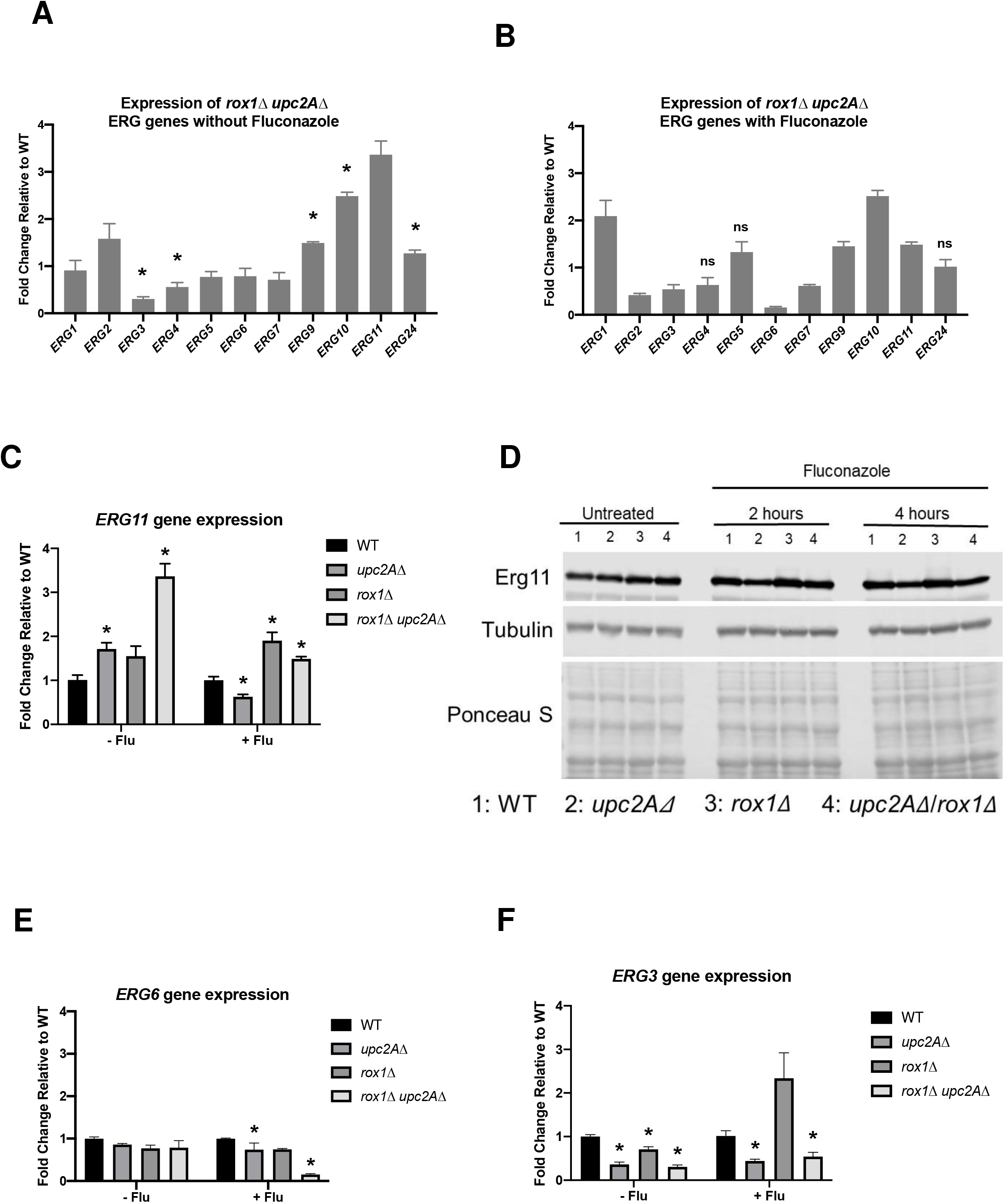
Deletion of r*ox1*Δ in *upc2A*Δ restores *ERG11* expression to near wild type levels in the presence of fluconazole. A. The expression of the indicated *ERG* genes was determined by quantitative RT-PCR and is normalized to WT. B. The expression of *ERG11* in the indicated strains. C. Western blot of the indicated strains grown in YPD or YPD + 8 µg/mL fluconazole for 4 hours. D. Expression of *ERG6* in the indicated strains. F. Expression of *ERG3* in the indicated strains. The strains were incubated in YPD or YPD+ 8 µg/mL fluconazole for 4 hr. The bars are means of three biological replicates performed in technical triplicate. The error bars indicate standard deviation and Student’s t test was used to determine statistical significance. Statistical significance relative wild type is indicated by an *.

Overall, 8 of the 11 *ERG* genes that we examined are expressed in the *rox1*Δ *upc2A*Δ mutant at levels that correspond to ≥ 75% of WT in the presence of fluconazole (Fig. 7A). Of these 8, *ERG11, ERG2* and *ERG3* are reduced by at least two-fold in the fluconazole-treated *upc2A*Δ mutant. Surprisingly, *ERG6* is not significantly reduced in either the *rox1*Δ or the *upc2A*Δ mutants in the presence of fluconazole (Fig. 7D). In the *rox1*Δ *upc2A*Δ mutant, however, *ERG6* expression is dramatically reduced, indicating that there is a strong negative genetic interaction between *upc2A*Δ and *rox1*Δ mutations with respect to *ERG6* expression. As discussed above, the reduced expression of *ERG6* in the fluconazole-treated *rox1*Δ *upc2A*Δ mutant is consistent with the sterol profiling data indicating decreased flux through that enzyme.

The expression of *ERG3* is also reduced in the fluconazole-treated *rox1*Δ *upc2A*Δ mutant (Fig. 7A) and correlates with sterol profiling evidence of reduced flux through that enzyme as well. *ERG3* is upregulated in the *rox1*Δ mutant in the presence of fluconazole and downregulated in the *upc2A*Δ mutant (Fig. 7E). In contrast to *ERG11*, however, the derepression of *ERG3* in *rox1*Δ does not compensate for its reduced expression in *upc2A*Δ and, indeed, its expression remains reduced in the *rox1*Δ *upc2A*Δ mutant (Fig. 7E). This provides further support for the conclusion that the relationship between Rox1 and Upc2A is more complex than a repressor and activator of a shared set of genes. However, the wild type levels of *ERG11* expression in the fluconazole-treated *rox1*Δ *upc2A*Δ mutant coupled with the reduced expression of *ERG3* and *ERG6* provide a potential mechanism for the ability of *rox1*Δ to suppress *upc2A*Δ fluconazole hyper-susceptibility.

### Transcriptional profiling of *upc2A*Δ, *rox1*Δ, and *upc2A*Δ *rox1*Δ strains

To characterize the effect of the double mutant on genome-wide gene expression, we performed RNA-Seq on *upc2A*Δ, *rox1*Δ, and *upc2A*Δ *rox1*Δ strains in the presence and absence of fluconazole. The processed data sets for all strains and conditions are provided in Supplementary Table 2. The role of *ROX1* in *C. glabrata* gene expression has not previously been characterized. As discussed above its role in the repression of hypoxia-related genes during normoxia is well-characterized in *S. cerevisiae* (15, 23). In log phase growth in YPD, 90 genes are upregulated by 2-fold with a corrected p value ≤ 0.05 (Fig. 8A). As shown in Fig. 8B, 5 genes involved in hypoxic response were upregulated, suggesting that Rox1 function is conserved between *C. glabrata* and *S. cerevisiae*. Previously reported transcriptional profiling indicated that ScRox1 suppresses cell wall-related genes (23). Consistent with those findings, cell wall genes were enriched (GO term, corrected p = 3.4 × 10^−4^; Fig. 8B), further supporting conservation of function between ScRox1 and Rox1.

**Figure 8.**
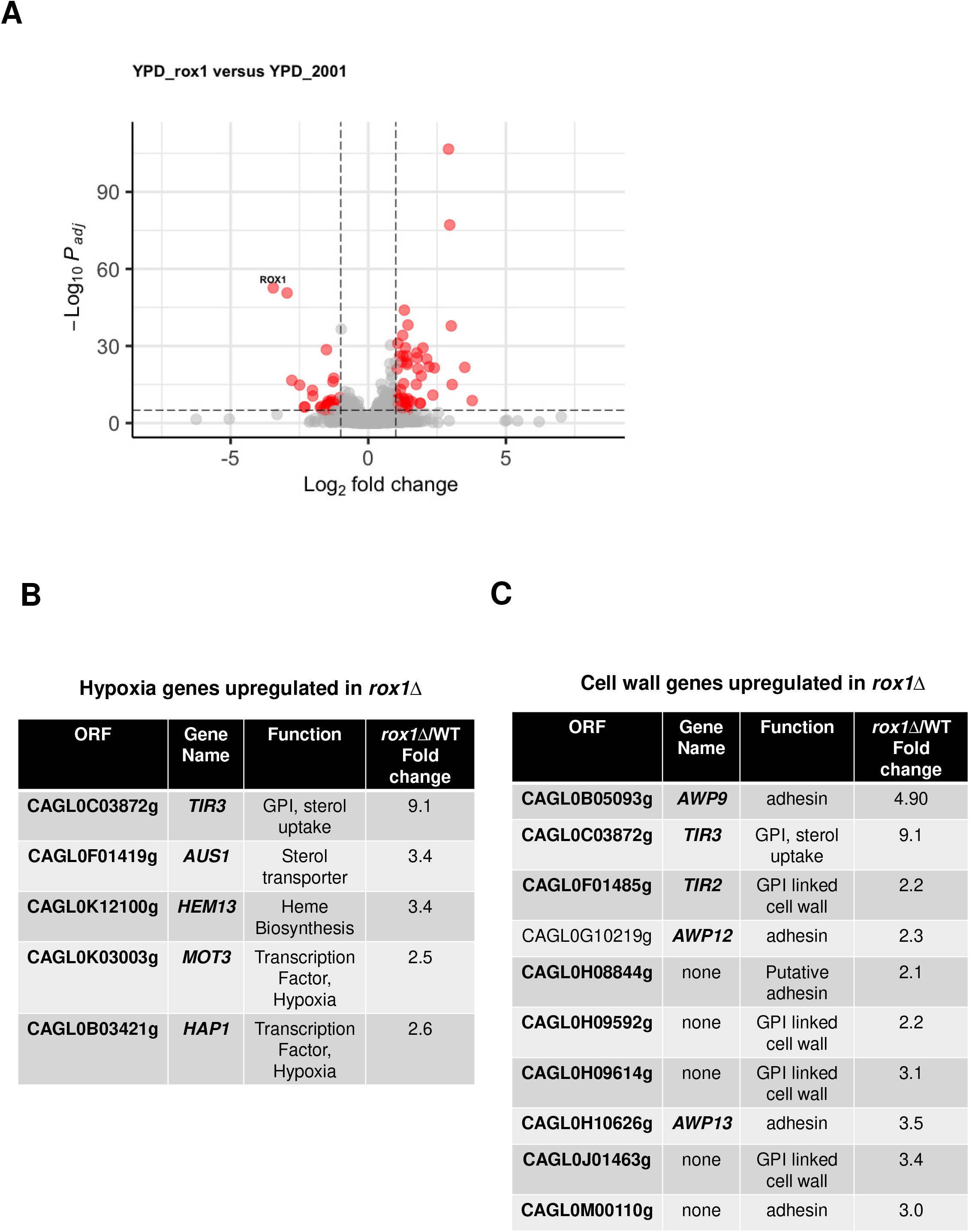
Rox1 represses the expression of hypoxia and cell wall-related genes under normoxia. A. Volcano plot comparing the expression of genes in *rox1*Δ to WT in log phase cultures incubated for 4 hours in YPD at 30°C. Red indicates genes that are changed by ±2-fold with a corrected p value < 0.05. The expression of hypoxia-related (B) and cell wall-related genes in *rox1*Δ mutant relative to WT under conditions indicated above.

In the presence of fluconazole, Rox1 would be expected to no longer repress genes required for the cellular response to sterol depletion and, thus, those genes may not be differentially regulated in the deletion mutant relative to WT. In the presence of fluconazole, 93 genes are upregulated in the *rox1*Δ mutant. Of the 93 genes, 48 remain upregulated in the presence of fluconazole while 42 are no longer upregulated; the latter set would be candidate genes that may be Rox1-regulated in the presence of sterol stress. This set of genes is enriched for carbohydrate metabolic processes (p = 7.7 × 10^−4^) and oxidation-reduction process (p = 3.1 × 10^−3^), two classes of genes also found to be Sc*rox1*-dependent during sterol stress (23). The genes upregulated only in the presence of fluconazole were not enriched for any process or cellular compartment by GO term analysis while those in common were enriched in cell wall-related genes (p = 9.25 × 10^−6^) and included *TIR3*, a cell wall gene implicated in sterol uptake (11, 26).

The effect of the *upc2A*Δ mutation on gene expression in the presence and absence of fluconazole has recently been reported by Vu et al (27) and our results are consistent with those data. We used our *upc2A*Δ data to compare to the *rox1*Δ *upc2A*Δ (Fig. 9A) in the presence of fluconazole to determine whether non-*ERG* genes that were downregulated in *upc2A*Δ were returned to WT levels of expression by the deletion of *ROX1*. Nineteen of the 32 genes downregulated in the fluconazole-treated *upc2A*Δ mutant was either equal to WT expression or upregulated in the *rox1*Δ *upc2A*Δ mutant. As expected, this set of genes was enriched for ergosterol biosynthesis genes (p = 5.8 × 10^−9^). In addition, UMP biosynthesis genes (p = 8.8 × 10^−8^) and oxidation-reduction genes (p = 2.8 × 10^−5^) were enriched, although it is difficult to connect these genes to fluconazole response. Thus, it appears that loss of Rox1 function restores the expression of some Upc2A dependent genes to levels comparable to WT cells in the presence of fluconazole.

**Fig. 9.**
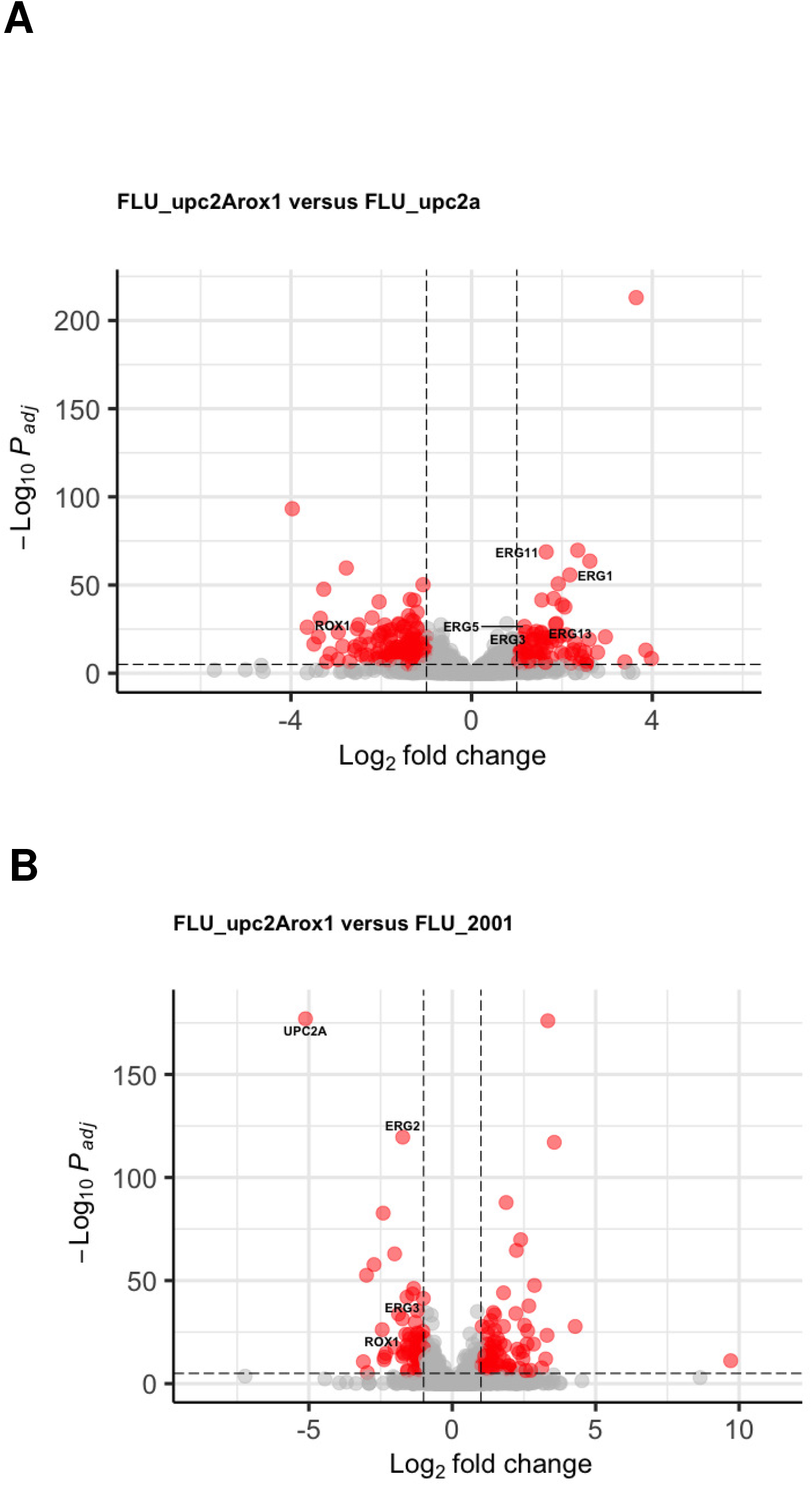
Transcriptional profile of *rox1*Δ *upc2A*Δ relative to WT and to the single mutant strains in the presence of fluconazole. The strains were incubated in YPD + 8 µg/mL fluconazole for 4 hours. A. Volcano plots comparing *rox1*Δ *upc2A*Δ to *upc2A*Δ (A) and WT (B). Red indicates genes that are changed by ±2-fold with a corrected p value < 0.05. *ERG* genes meeting these criteria are labeled.

Relative to wild type in fluconazole, the *rox1*Δ *upc2A*Δ mutant has 158 genes that are upregulated. Both *upc2A*Δ and *rox1*Δ also have significant number of genes upregulated in the presence of fluconazole, 153 and 93 respectively. It seems likely that many of the genes that are upregulated in fluconazole-treated *upc2A*Δ are related to a compensatory response to the severe sterol stress of those cells. Consistent with that hypothesis, 15 cell wall genes are upregulated in the fluconazole-treated *upc2A*Δ cells (p = 0.04). These include the cell wall integrity pathway (CWIP) MAPK Slt2, a gene known to be induced and activated by sterol stress (28), along with *FKS2*, the gene coding a catalytic subunit of 1, 3-β-glucan synthase and a gene upregulated by activation of Slt2. Of the 153 genes that are upregulated in the *upc2A*Δ mutant, only 40 (26%) are among the 158 genes that are upregulated in the double mutant in the presence of fluconazole. This set is no longer enriched for cell wall genes and, indeed, none of the 15 cell wall genes remain upregulated. These observations are consistent with the hypothesis that the large set of genes upregulated in *upc2A*Δ represents an exaggerated compensatory response to sterol stress that is then largely eliminated by the deletion of *ROX1*.

Overall, our transcriptional profiling supports the conclusion that Rox1 function is reasonably well conserved between *S. cerevisiae* and *C. glabrata*. Deletion of Rox1 suppresses the *upc2A*Δ fluconazole hypersusceptibility phenotype, at least in part, by de-repressing *ERG* gene expression through a Upc2A-independent pathway. Finally, fluconazole-treated *upc2A*Δ cells display a compensatory transcriptional response that is no longer induced in the fluconazole-treated *rox1*Δ *upc2A*Δ double mutant.

## Discussion

The ergosterol biosynthesis pathway is not only an essential component of fungal membrane biology but also the target of two of the three classes of drugs that are used to treat human fungal infections. Despite its centrality to fungal biology and antifungal drug therapy, our models for the regulation of ergosterol biosynthesis are relatively limited, particularly in the case of pathogenic fungi (13). In general, current understanding of the transcriptional regulation of *ERG* gene expression is limited almost entirely to the role of SREB transcription factors or functionally homologous, Zn cluster transcription factors such as Upc2A (14). Although many *ERG* genes are known to be essential, none of the TFs that regulate their expression are essential, indicating that additional factors are likely to contribute to the regulation of *ERG* genes. In addition, over-expression experiments in *S. cerevisiae* have shown that inappropriately high expression of *ERG* genes reduces fitness likely due to the fact that sterols can be toxic to the cell (29). As such, it seemed likely that negative regulators of sterol biosynthesis may be required to maintain sterol homeostasis. Our screen was designed to identify either gain-of-function mutants in positive regulators of *ERG* gene expression or loss-of-function mutants in negative regulators and yielded two negative regulators of *ERG* gene expression: Rox1 and potentially Cst6.

Based on previous work regarding the roles of ScRox1 and ScUpc2/Ecm22 in the regulation of hypoxia and sterol stress, the ability of *rox1*Δ to suppress the *upc2A*Δ phenotype is somewhat unexpected. Specifically, in *S. cerevisiae*, ScRox1 represses many genes that are activated by ScUpc2 and, thus, loss of the Rox1 repressor should require the presence of the cognate activator for shared genes to be expressed (23). In the presence of fluconazole, the deletion of *ROX1* in the *upc2A*Δ leads to normalization of some, but not all, *ERG* gene expression and increased overall levels of ergosterol relative to WT *C. glabrata* cells. Accordingly, there must be a transcription factor in addition to Upc2A that is able to activate the expression of at least some *ERG* genes in the presence of fluconazole; otherwise, loss of the repressor Rox1 would not be able to restore growth during sterol stress because *UPC2A* was deleted in the parental strains used in the screen. This factor also must be sufficient to restore growth under hypoxic conditions to *upc2A*Δ mutants because the *rox1*Δ mutation suppresses the hypersensitivity of *upc2A*Δ to those conditions.

Upc2A is currently the only factor that has been directly implicated in the regulation of *ERG* genes in *C. glabrata* (9-14). Although Upc2B would seem a logical candidate for this factor, it has not been shown to play a role in the regulation of *ERG* genes (10). In the fluconazole-treated *upc2A*Δ *rox1*Δ mutants, *UPC2B* expression is modestly reduced (Table S2). This is consistent with previously published data indicating that Upc2A is required for induction of *UPC2B* expression (10). We cannot completely rule out the possibility that Upc2B contributes to the regulation of *ERG* gene expression in the absence of *UPC2A* but the current data seem more consistent with an alternative explanation.

A second candidate factor is Hap1 which, based on the large-scale genetic phenotyping project, is required for WT growth in fluconazole (17). *HAP1* is upregulated in the *rox1*Δ mutant in the absence of fluconazole and is expressed near WT levels in the presence of fluconazole in the *upc2A*Δ *rox1*Δ double mutant (Table S2). Hap1 is also involved in the expression of *ERG* genes under unstressed conditions in *S. cerevisiae* (30). Certainly, other factors with less obvious effects on fluconazole stress could also be involved.

Another question raised by our observations is, why isn’t the putative missing *ERG*-gene regulating factor activated in the presence of fluconazole to compensate for the *upc2A*Δ mutationã The genetic interaction of *upc2A*Δ and *rox1*Δ imply that the presence of Rox1 prevents this factor from positively regulating *ERG* gene expression in the absence of Upc2A. The scheme in Fig. 10 outlines a genetic circuitry that is consistent with our data. The data also suggest that Rox1 may still repress the expression of some genes during treatment with fluconazole or under low oxygen conditions. However, the only transcription factor that is upregulated in the *rox1*Δ *upc2A*Δ mutant is Stb5 which has been shown as a negative regulator of azole susceptibility by decreasing expression of *CDR1* and other efflux pumps (31). Another possibility is that Upc2A may be needed to fully displace Rox1 from promoters and Rox1, in turn, may directly interfere with the binding of this other positive activator to *ERG* gene promoters. Additional work will be needed to identify the missing positive regulator and understand the mechanism for the interactions between the positive and negative regulators of ergosterol biosynthesis.

**Fig. 10.**
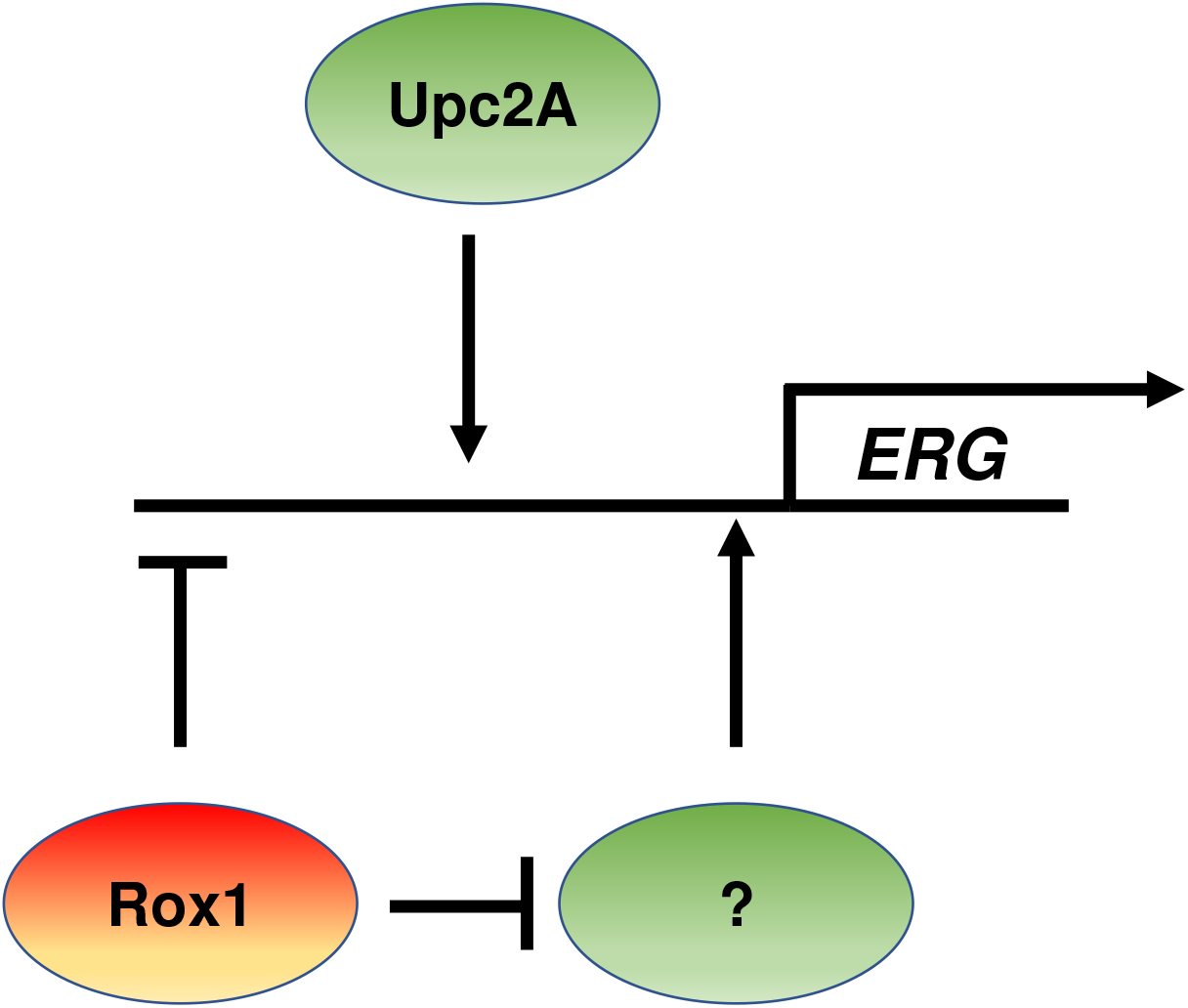
Schematic for proposed circuit with unidentified positive regulator of *ERG* gene expression implicated in this work.

The biochemical mechanism by which the *rox1*Δ mutation suppresses *upc2A*Δ fluconazole hyper-susceptibility is by restoring *ERG11* and other *ERG* genes to wild type or near wild type levels in the presence of fluconazole and, thereby, leading to increased ergosterol content in *rox1*Δ *upc2A*Δ cells relative to *upc2A*Δ. The sterol profile of the fluconazole-treated double mutant differs from WT indicating that normal flux through the ergosterol pathway has not been re-established. Specifically, sterols that are dependent on Erg3 and Erg6 are reduced while the sterols that serve as Erg3 and Erg6 substrates are increased. This correlates with reduced expression of the *ERG3* and *ERG6* genes in the double mutant. The fact that the double mutant has *ERG11* expression restored to WT levels but not *ERG3* and *ERG6* further emphasizes the complexity of ergosterol biosynthesis gene regulation and that Upc2A and Rox1 are not simply on/off switches for a common set of genes.

Furthermore, this apparent reduction in Erg3 and Erg6 leads to a reduction in the toxic sterol, 14α methyl-ergosta-8,24(28)-dien-3β, 6α-diol, that accumulates in cells treated with the Erg11-inhibiting azoles (25). Thus, the *rox1*Δ *upc2A*Δ strain has both an increase in ergosterol and a decrease in the toxic sterol relative to *upc2A*Δ and wild type cells. This not only provides a mechanism for the restoration of wild type azole susceptibility through deletion of *ROX1* in the *upc2A*Δ mutant, but also indicates that changes in the relative expression of *ERG* genes can lead to profound changes in azole susceptibility. In principle, these data suggest that SNPs in the promoter regions of *ERG* genes or epigenetic alterations in gene expression could lead to changes in azole susceptibility without any change to protein coding sequences.

Lastly, we consider the implications of our findings in *C. glabrata* to *C. albicans* which, as noted in the discussion, has a number of distinctions with respect to the genetic regulation of fluconazole susceptibility. Most notably, the Rox1 homolog in *C. albicans* is Rfg1, and, although it is an HMG class repressor, it has no role in the suppression of hypoxia genes. Instead, it represses filamentation (32, 33). Although a Upc2A homolog is present and has been shown to function very similarly, there are no Upc2B/ScEcm33 homologs in *C. albicans*. Taken together, our data provide strong evidence for the conclusion that, although the ergosterol biosynthesis pathway is highly conserved among pathogenic and model yeast, significant changes have occurred in terms of its transcriptional regulation. A more detailed understanding of these distinctions may provide additional insights to the possible mechanisms of resistance to inhibitors of ergosterol biosynthesis.

In summary, our genetic suppressor study provides new insights into the regulation of ergosterol biosynthesis and fluconazole response pathways in *C. glabrata*. These data highlight the complex nature of the transcriptional pathways that maintain ergosterol homeostasis during normal growth and under sterol stress. Future studies will further explore mechanisms by which the putative Rox1 suppressor pathway is regulated and how Cst6 affects ergosterol biosynthesis in *C. glabrata*.

## Materials and methods

### Media, reagents and general methods

Yeast peptone dextrose (YPD), synthetic complete medium with amino acid dropout/supplements were prepared using standard recipes (34) for both liquid and plate (2% agar) based media. Strains were streaked from frozen stocks on to YPD plates and incubated at 30°C. For all experiments, strains were pre-cultured in liquid medium with shaking at 30°C overnight. Fluconazole was obtained as powder from Sigma.

### Strain and Plasmid construction

The parental *upc2A*Δ strain for the laboratory evolution experiment was constructed in the BG2 background (17) as described in Vu et al. using the recyclable NAT cassette flanked by 1kb of homology to the upstream and downstream regions of *UPC2A* (18). Correct integration was confirmed by PCR as was the absence of the *UPC2A* ORF. The *rox1*Δ *upc2A*Δ mutant was constructed in a similar manner in KK2001 background by deleting the *ROX1* coding region from the previously reported *upc2A*Δ *ERG11*-HA strain (27).

### *C. glabrata* transformation

Transformations of *C. glabrata* were performed using the lithium acetate method (18). Post heat shock, cells were grown overnight at 30°C at 200 rpm for drug selection. Overnight cultures were spun down and plated on YPD plates supplemented with 100 µg/ml of nourseothricin (NAT) (Jena Bioscience, Jena, Germany). Colonies were purified by plating on YPD plates containing 200 µg/ml NAT.

### Experimental evolution of suppressors of *upc2A*Δ fluconazole hyper-susceptibility

Three independent colonies of *upc2A*Δ mutant in the His-BG2 background precultured overnight in YPD. The three culture was used to inoculate three separate 96-well plates at 1000 CFU/well in YPD containing 0.25 µg/mL fluconazole (1/4 MIC for this strain). The plates were incubated at 30°C until the wells were saturated. The cultures were serially passaged by transferring 2 µL of each well to a fresh plate containing 2X the fluconazole concentration as the previous plate up to a final concentration of 64 µg/mL. Surviving lineages were plated on YPD and random colonies from each were serially streaked on YPD without fluconazole. The ability of the strains to grow at 64 µg/mL fluconazole was re-tested. From this experiment, 12 independent strains derived from *upc2A*Δ with the ability to grow at 64 µg/mL were isolated: See table if Fig. 2A for names of the resistant strans. The genomes of the strains were sequenced using Illumina at the University of Iowa Genomics Core. The genomes were assembled and SNPs called using the Lasergene platform. The sequencing data are deposited.

### Spot dilution assay

Strains were grown overnight at 30°C at 200 rpm in liquid YPD, SCM or SCM without methionine and cysteine. For YPD and YPD + fluconazole, the strains were diluted to OD_600_ 1 and plated with 10-fold serial dilutions. Plates were incubated at 30°C or 37°C for 1-3 days before photographing them.

### Hypoxia growth assay

Cells were grown in liquid YPD overnight at 30°C at 200 rpm and diluted to (OD) of 1. Cells were plated on YPD and serially diluted 10-fold for spot dilutions. Plates were sealed in BD GasPak EZ Anaerobe Gas Generating Pouch System with Indicator and placed in 30°C incubator or placed directly in incubator. Plates were captured after 48 hours.

### Western blot analysis

Strains were incubated in YPD or with YPD in fluconazole for 4 hours. Proteins were extracted as previously described (Shahi *et al*., 2010), resuspended in 120 μl of urea-SDS sample buffer (8 M urea, 5% SDS, 1% 2-mercapto-ethanol, 40 mM Tris-HCl pH 8.0, bromophenol blue) and incubated at 37° C for 3 hours by occasional vortexing. Aliquots of 12 μl were resolved on precast ExpressPlus 4-12% gradient gel (GenScript #M41212). Proteins were electroblotted to nitrocellulose membrane, blocked with 5% nonfat dry milk. Membrane was probed with polyclonal anti-Cdr1 antibody (27) in the final dilution of 1:1000 overnight at 4° C and with 12G10 anti-alpha-tubulin monoclonal antibody (Developmental Studies Hybridoma Bank at the University of Iowa) in the final dilution of 1:5000 for 15 minutes at room temperature. Secondary Li-Cor antibodies IRD dye 680RD goat anti-rabbit (#926-68021) and IRD dye 800LT goat anti-mouse (#926-32210) were used in the final dilution of 1:10 000. Li-Cor infrared imaging system (Li-Cor, application software version 3.0) was used to detect the signal. The data represent results of two biologically independent experiments.

### Sterol profiling

Overnight cultures from single colonies of *C. glabrata* strains were used to inoculate 20 mL YPD (starting OD_600nm_ 0.20) in the absence (DMSO control, 1% v/v) or presence of 8 µg/mL fluconazole (stock prepared in DMSO, final concentration 1% v/v DMSO). Cultures were grown at 30°C for 16 hr at 180 rpm. Cells were then pelleted and washed with ddH_2_O before splitting each sample for sterol extraction and dry weight determination. Sterols were extracted and derivatized as previously described (35). An internal standard of 5 µg of cholesterol was added to each sample and lipids were saponified using alcoholic KOH and non-saponifiable lipids extracted with hexane. Samples were dried in a vacuum centrifuge and were derivatized by the addition of 0.1 mL BSTFA TMCS (99:1, Sigma) and 0.3 mL anhydrous pyridine (Sigma) and heating at 80°C for 2 hours. TMS-derivatised sterols were analysed and identified using GC/MS (Thermo 1300 GC coupled to a Thermo ISQ mass spectrometer, Thermo Scientific) and Xcalibur software (Thermo Scientific). The retention times and fragmentation spectra for known standards were used to identify sterols. Integrated peak areas were determined to calculate the percentage of total sterols. Ergosterol quantities were determined using standard curves of peak areas of known quantities of cholesterol and ergosterol. Sterol composition and ergosterol quantities were calculated as the mean of three replicates. The statistical significance of the differences between strains was determined using the means and standard error of the means and Student’s t test with p < 0.05 indicating statistical significance. The complete sterol data and summary are provided in Supplementary Table 1.

### Transcriptional analysis by quantitative reverse transcription-PCR (qPCR)

Strains were incubated overnight in liquid YPD at 30°C at 200 rpm, back diluted into fresh YPD and grown to mid-log phase. Cells were harvested at mid-log phase and resuspended into fresh YPD or YPD with 8 µg/ml of fluconazole. Cultures were incubated for 4 hours, harvested and total RNA was isolated with MasterPure™ Yeast RNA Purification Kit. The RNA was reverse transcribed using iScript cDNA synthesis kit (170-8891; Bio-Rad) and qPCR reaction performed using IQ SyberGreen supermix (170-8882; Bio-Rad). *ACT1* expression was used as the normalization standard and relative expression between strains and conditions was determined using the ΔΔ_CT_ method. Experiments were performed in biological duplicate with technical triplicates and the statistical significance was determined using Student’s t test with significance limit of p < 0.05.

### RNA sequencing methods

The strains were grown with and without fluconazole as described for single gene expression analysis by qRT-PCR. RNA sequencing performed using Illumina MiSeq for stranded mRNA. Libraries were prepared with paired end adapters using Illumina chemistries per manufacturer’s instructions, and sequencing of libraries were performed with read lengths of approximately 150bp with at least 50 million reads per sample. RNA sequencing reads were imported into CLC genomics workbench 20.0. Sequences were trimmed and aligned to reference sequence (https://www.ncbi.nlm.nih.gov/assembly/GCF_000002545.3) ASM254v2_genomic-1 with paired reads counted as one. Mismatch, insertion, and deletion costs were set to CLC default parameters. Differential expression analysis for whole transcriptome RNA-seq was performed using CLC genomics workbench. Wald test was used in all group pairs against the indicated control group. Statistics based on the fit of a generalized linear model with a negative binomial distribution. *C. glabrata* gene annotations were obtained from Candida Genome Database. The fold changes and corrected p-value for all genes and all conditions are provided in Supplementary Table 2.

## Acknowledgements

This work was supported by NIH grants 5R01AI52494 (SMR) and 7R01AI131620 (PDR). the funders had no role in study design, data collection and interpretation, or the decision to submit the work for publication

